# Transcripts and genomic intervals associated with variation in metabolite abundance in maize leaves under field conditions

**DOI:** 10.1101/2024.08.26.609532

**Authors:** Ramesh Kanna Mathivanan, Connor Pederson, Jonathan Turkus, Nikee Shrestha, J. Vladimir Torres-Rodriguez, Ravi V. Mural, Toshihiro Obata, James C. Schnable

## Abstract

Plants exhibit extensive environment-dependent intraspecific metabolic variation, which likely plays a role in determining variation in whole plant phenotypes. However, much of the work seeking to use natural variation to link genes and transcript’s impacts on plant metabolism has employed data from controlled environments. Here we generate and employ data on variation in the abundance of twenty-six metabolites across 660 maize inbred lines under field conditions. We employ these data and previously published transcript and whole plant phenotype data reported for the same field experiment to identify both genomic intervals (through genome-wide association studies) and transcripts (through both transcriptome-wide association studies and an explainable AI approach based on the random forest) associated with variation in metabolite abundance. Both genome-wide association and random forest-based methods identified substantial numbers of significant associations including genes with plausible links to the metabolites they are associated with. In contrast, the transcriptome-wide association identified only six significant associations. In three cases, genetic markers associated with metabolic variation in our study colocalized with markers linked to variation in non-metabolic traits scored in the same experiment. We speculate that the poor performance of transcriptome-wide association studies in identifying transcript-metabolite associations may reflect a high prevalence of non-linear interactions between transcripts and metabolites and/or a bias towards rare transcripts playing a large role in determining intraspecific metabolic variation.

## Introduction

Plants can produce a wide array of metabolites with diverse structures that perform essential roles in growth, cellular regeneration, resource allocation, development, and responses to biotic and abiotic stresses. While the total number of metabolites produced by plants likely exceeds one million, each plant species typically synthesizes between several thousand and several tens of thousands^1, 2^. In addition to interspecies metabolic diversity, substantial metabolic diversity exists between members of the same species (intraspecies diversity). Investigating the genetic determinants of intraspecies variation in plant metabolism can provide insight into both the enzymes responsible for specific steps in metabolic pathways and also the role variation in plant metabolism plays in determining whole plant phenotypes^2, 3^. Variation in the abundance of lignin precursors is correlated with variation in biomass production in arabidopsis and maize^4, 5^. Many metabolic and morphological trait pairs exhibited significant correlations in an analysis of 64 metabolite traits and 35 morphological traits scored across a tomato mapping population^6^. QTL for nine of twenty-six whole plant phenotypes evaluated in potato colocalized with QTL for variation in the abundance of one or more of 85 metabolites profiled in the same population^7^.

Plant metabolic traits tend to be moderately heritable within species. More than half of metabolic features profiled in a rice diversity panel exhibited heritability coefficients (H^2^) generally greater than 0.5 and for nearly one-quarter, H^2^ exceeded 0.7^8^. Metabolic quantification of a population of 289 maize genotypes identified 26 metabolites where one or more genetic markers were significantly associated with variance in abundance in the leaves of maize seedlings grown in controlled environment conditions including a locus on chromosome 9 associated with variation in lignin precursors^5^. More than 1,400 genetic loci were significantly associated with variation in the abundance of 983 metabolites measured in mature (dry) maize kernels^9^. Comparative GWAS for the abundance of metabolites in dried seeds conducted in rice and maize identified 420 and 292 loci associated with 123 metabolites in the two species, respectively. These hits included 42 associated with homologous loci in both species^10^.

A combination of genome-wide association studies and transcriptome-wide association studies was able to link thirteen genes to variation in the abundance of tocochromanol (vitamin E) including five genes not previously linked to this metabolite^11^. Transcriptome-wide association studies can provide advantages in interpretation as they appear to be less influenced by linkage disequilibrium and more frequently identify a single candidate gene per genomic interval than genome-wide association studies^12^. Combined genome-wide and transcriptome-wide association studies have also been used to identify loci associated with variation in seed oil content in *Brassica napus*^13^. This frequent use of seeds for population-level metabolic profiling^9–11, 13^ is likely due to the significant economic and food security importance of crop seeds, as well as the practical challenges associated with collecting and sampling equivalent vegetative tissues from large populations grown under field-relevant conditions

Here we seek to identify loci associated with variation in metabolite abundance under field conditions in mature leaf tissue employing data generated from the Wisconsin Diversity panel, a large maize association panel selected to capture the genetic and phenotypic diversity present among the set of maize genotypes with the potential to complete their life cycle in temperate North America^14^. This panel has been previously resequenced, providing an extremely high marker density for genome-wide association studies^15^, and we leveraged a parallel RNA-seq experiment with profiled gene expression using leaf samples collected from the same plants on the same day as those employed for quantification of metabolites^16^. We were able to identify a total of 240 genes associated with metabolite variations and one gene that was identified between TWAS and RF. This study not only highlighted genes directly linked to metabolite production, such as N-acetyl-gamma-glutamyl-phosphate reductase with L-glutamic acid but also revealed other genes crucial for various aspects of plant metabolism, especially resistance against biotic and abiotic stresses. Additionally, we identified three loci associated with variation in both metabolic and non-metabolic traits. The data and analyses presented here will allow future research to functionally validate links between genes and metabolites as well as metabolites and whole plant phenotypes.

## Results

A set of 26 unique metabolites was successfully quantified via GC-MS across mature leaf tissue samples collected from 660 maize inbred lines. A total of 660 unique sets of measurements were generated, representing different genotypes. The dataset initially included 707 samples, of which 47 are biological replicates from tissue samples collected from different plants of the same genotype within the same experiment. Additionally, 88 of these samples are duplicates of the replicated check, B97, which were not included in the count of unique genotypes, resulting in 660 unique sets of measurements. Several metabolites exhibited high biological repeatability across these replicates. Threonine showed the highest repeatability (r =0.87), suggesting strong genetic control. Other metabolites such as raffinose (r=0.82), chlorogenic acid (r=0.81), malic acid (r=0.80), and galactonic acid (r=0.77) also displayed repeatability values indicative of reasonable heritability. The average biological repeatability was 0.57 (Figure 1A, S1). These values were compared to average biological repeatability of 0.75 for 28 whole plant traits^17^, 0.42 for 10 hyperspectral traits^18^, and 0.25 for 3 photosynthetic traits scored for the same maize genotypes within the same experiment (Figures S2, S3 and S4).

**Figure 1.**
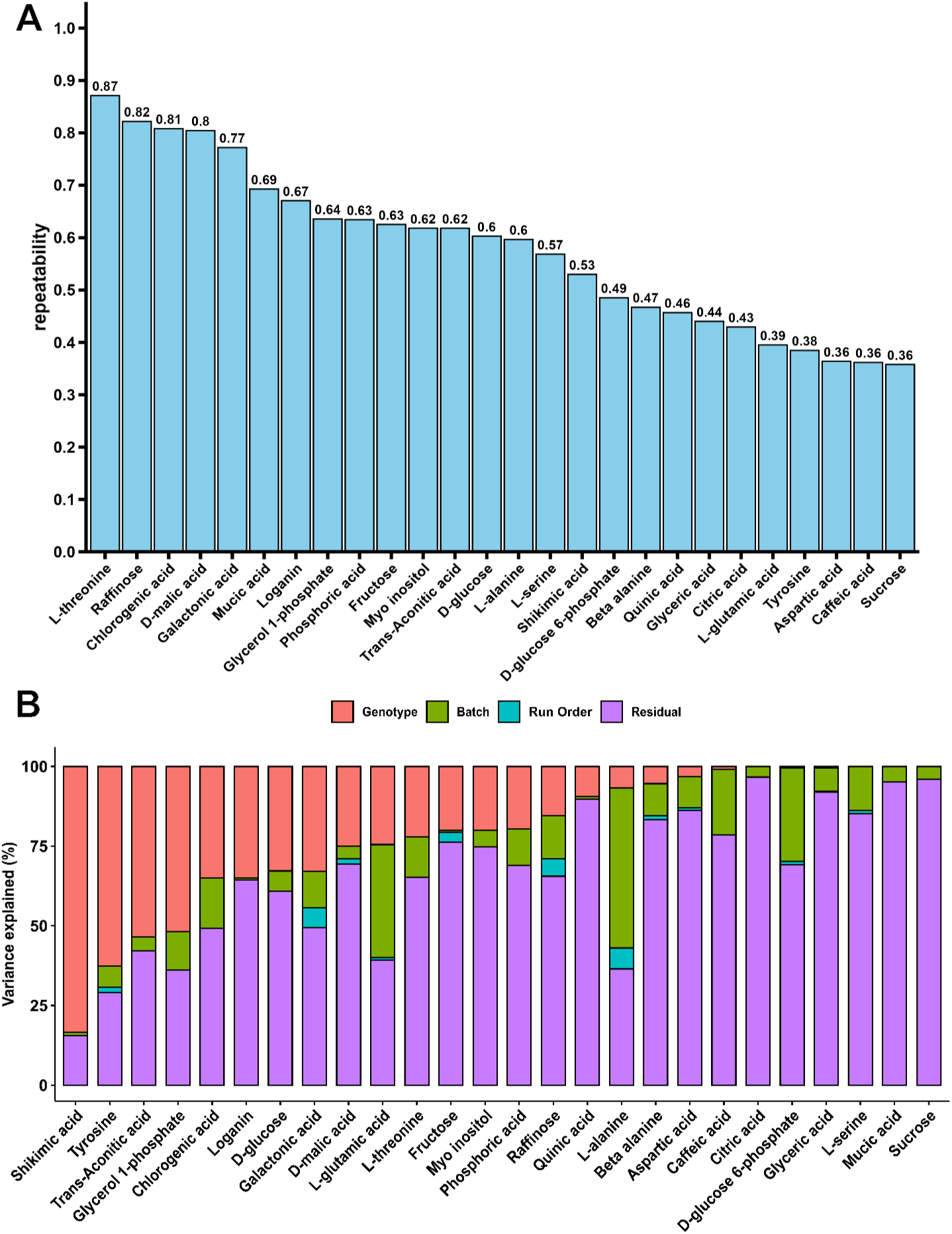
Factors explaining variation in metabolite abundance across a maize diversity panel. **A)** Estimated repeatability of measured metabolite abundance for each metabolite quantified in this study. Repeatability is defined as the proportion of total variance in metabolite abundance which can be explained by genotype in a dataset of 47 maize genotypes sampled and analyzed twice independently from different plants in the same field. **B)** Proportion of total variance in each metabolite’s abundance explained by the factors genotype, batch, run order, and the remaining residual variance. Genetic variance in panel B is not equivalent to repeatability in panel A as a model with more factors was fit to explain variance in panel B.

We did not observe strong evidence of clustering among different groups of samples (Figure S5), However, a significant proportion of the non-genetic variance for some individual metabolites could be explained by variation between different batches of samples, which refers to groups of samples processed at different times or differences in data generated and analyzed earlier or later within a given batch during the GC-MS run (Figure 1B). Several expected correlations were observed between different metabolites profiled, such as the positive correlation between glucose and fructose, both of which are sugars commonly involved in similar metabolic pathways, and between quinic acid and shikimic acid, which are both intermediates in the same biosynthetic pathway (Figure S6). The abundance of a number of metabolites was also significantly correlated with whole plant phenotypes scored in the same field (Figure S7). Shikimic acid abundance measured in the field at the late vegetative/early flowering stage showed a significant positive correlation with plant height (r = 0.23; p < 0.0001) and a significant negative correlation with percent ear fill (r = −0.25; p < 0.0001) at harvest. Similarly, Beta-alanine abundance showed a significant positive correlation with 100 kernel mass (r = 0.22; p < 0.0001) at harvest.

After controlling for batch effects and order of quantification effects, 150 genetic markers were significantly associated with one or more of 26 metabolites at a resampling model inclusion probability (RMIP) threshold ≥ 0.1 (Figure 2). Among these 150 genetic markers, five were found to be associated with two different metabolites. The 17 most supported metabolite-genetic marker associations all exceeded RMIP ≥ 0.3 (Table 1) and included two markers each associated with variation in the abundance of phosphoric acid, chlorogenic acid, galactonic acid, and trans-aconitic acid, and one genetic mark each associated with variation in the abundance of glyceric acid, shikimic acid, L-serine, quinic acid, raffinose, sucrose, tyrosine, D-glucose, and fructose. In seven cases a gene known to play a specific role in plant metabolism was located within 50 kilobases of a genetic marker linked to metabolic variation at RMIP ≥ 0.3 and markers within the gene appeared to be in significant linkage disequilibrium with the genetic marker identified via GWAS (Figure S8;Table 1). However, in many cases, the windows around individual metabolite abundance-associated genetic markers, defined by linkage disequilibrium, included multiple gene models, with a median of 4 gene models and a mean of approximately 5 gene models per interval.

**Figure 2.**
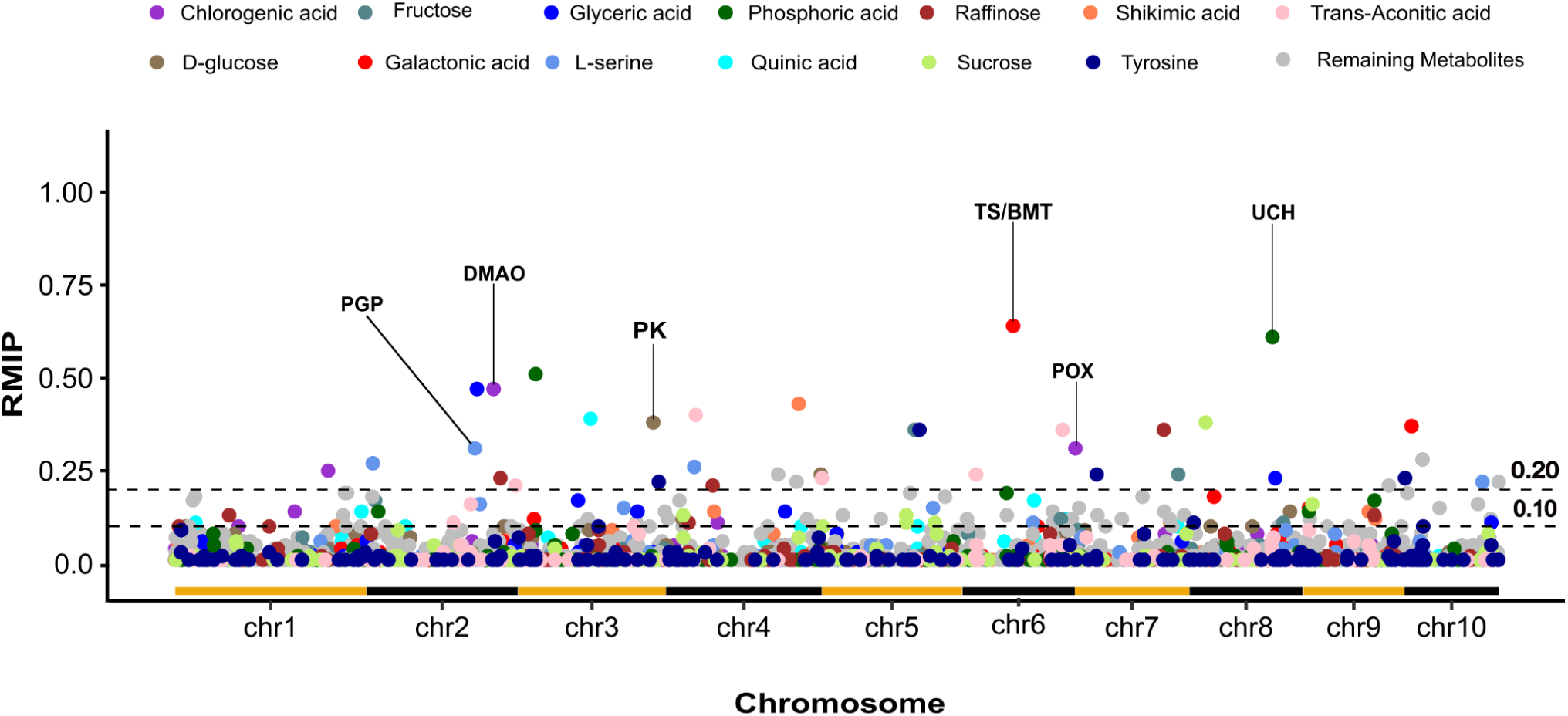
Genetic markers associated with metabolite variation via resampling model inclusion probability genome-wide association. Each circle’s position in the x-axis indicates the position of a given genetic marker on the maize genome, and its position on the y-axis indicates the proportion of resampling runs in which the marker was significantly associated with variation in the metabolite of interest via FarmCPU GWAS. For metabolites where at least one marker was associated with RMIP ≥ 0.3, color indicates the specific metabolite a given marker is associated with. Markers associated with all other metabolites tested are shown in gray. The two horizontal dashed lines mark RMIP = 0.2 and RMIP = 0.1. Text labels indicate genes discussed in the next near metabolite-associated markers. Alternating color horizontal lines along the x-axis indicate the start and end of each maize chromosome.

**Table 1.** Location, support, and closest gene model for each RMIP GWAS hit ≥0.3 shown in Figure 2. * In seven cases, a gene known to play a specific role in plant metabolism was located within 50 kilobases of a genetic marker linked to metabolic variation.

| Metabolites | RMIP | SNP | Candidate Gene | Distance from marker | Gene description |
| --- | --- | --- | --- | --- | --- |
| Galactonic acid | 0.64 | chr6_81874806 | Zm00001eb270570* | 38,129 | Theobromine synthase (TS) |
| Galactonic acid | 0.64 | chr6_81874806 | Zm00001eb270580* | 39,003 | Benzoate O-methyltransferase (BMT) |
| Phosphoric acid | 0.61 | chr8_133024421 | Zm00001eb354560* | 32,734 | Ubiquitin carboxyl terminal hydro-lase (UCH) |
| Phosphoric acid | 0.51 | chr3_28733643 | Zm00001eb126310 | 761 | NA |
| Glyceric acid | 0.47 | chr2_177080341 | Zm00001eb097540 | 29,285 | Endonuclease |
| Chlorogenic acid | 0.47 | chr2_203757097 | Zm00001eb104230* | 9,983 | Phosphoglycolate phosphatase (PGP) |
| Shikimic acid | 0.43 | chr4_212836690 | Zm00001eb201130 | 32,290 | NA |
| Trans-Aconitic acid | 0.40 | chr4_48138869 | Zm00001eb175400 | 8,808 | Uridyltransferase |
| Quinic acid | 0.39 | chr3_116669684 | Zm00001eb135350 | 53271 | NA |
| Sucrose | 0.38 | chr8_26033461 | Zm00001eb338670 | 14,868 | PWWP domain |
| D-Glucose | 0.38 | chr3_216820299 | Zm00001eb157570* | 19,238 | Protein kinase domain (PK) |
| Galactonic acid | 0.37 | chr10_11900210 | Zm00001eb408510 | 81,008 | PPR repeat |
| Raffinose | 0.36 | chr7_142676377 | Zm00001eb317720 | 10,013 | Proteasome subunit beta type-1 |
| Fructose | 0.36 | chr5_149005008 | NA | NA | NA |
| Tyrosine | 0.36 | chr5_156613475 | Zm00001eb239880 | 44,418 | NA |
| Trans-Aconitic acid | 0.36 | chr6_161018577 | Zm00001eb289030 | 34,988 | A/G-specific adenine glycosylase |
| Chlorogenic acid | 0.31 | chr7_1204204 | Zm00001eb298230* | 22,715 | Peroxidase |
| L-serine | 0.31 | chr2_173821786 | Zm00001eb096820* | 53,115 | Dimethylaniline monooxygenase (DMAO) |

Unlike genome-wide association studies, transcriptome-wide studies can frequently provide single gene resolution^12^, ameliorating the challenge of translating metabolite abundance associations to individual candidate genes. A previously published gene expression dataset generated from leaf tissue collected from the same plants at the same time as the leaf tissue samples employed for quantifying the abundance of metabolites^16^ was utilized to conduct transcriptome-wide association studies (TWAS) for metabolite abundance. The abundance of only three of the 26 metabolites was significantly associated with the expression of individual genes in the same leaves at a Bonferroni-corrected significance threshold of 0.05 (Figure 3;Figure S9). These included four genes whose transcript abundance was significantly associated with variation in the abundance of glycerol 1-phosphate, and one gene each associated with variation in L-glutamic acid and quinic acid. Variation in the expression of the same gene Zm00001eb431150, which encodes a Cu(2+)-exporting ATPase (CUEA), was linked to both variations in the abundance of glycerol 1-phosphate and L-glutamic acid. The sole gene whose transcript abundance was associated with quinic acid was Zm00001eb147850, which is a multi-copper oxidase (MCO).

**Figure 3.**
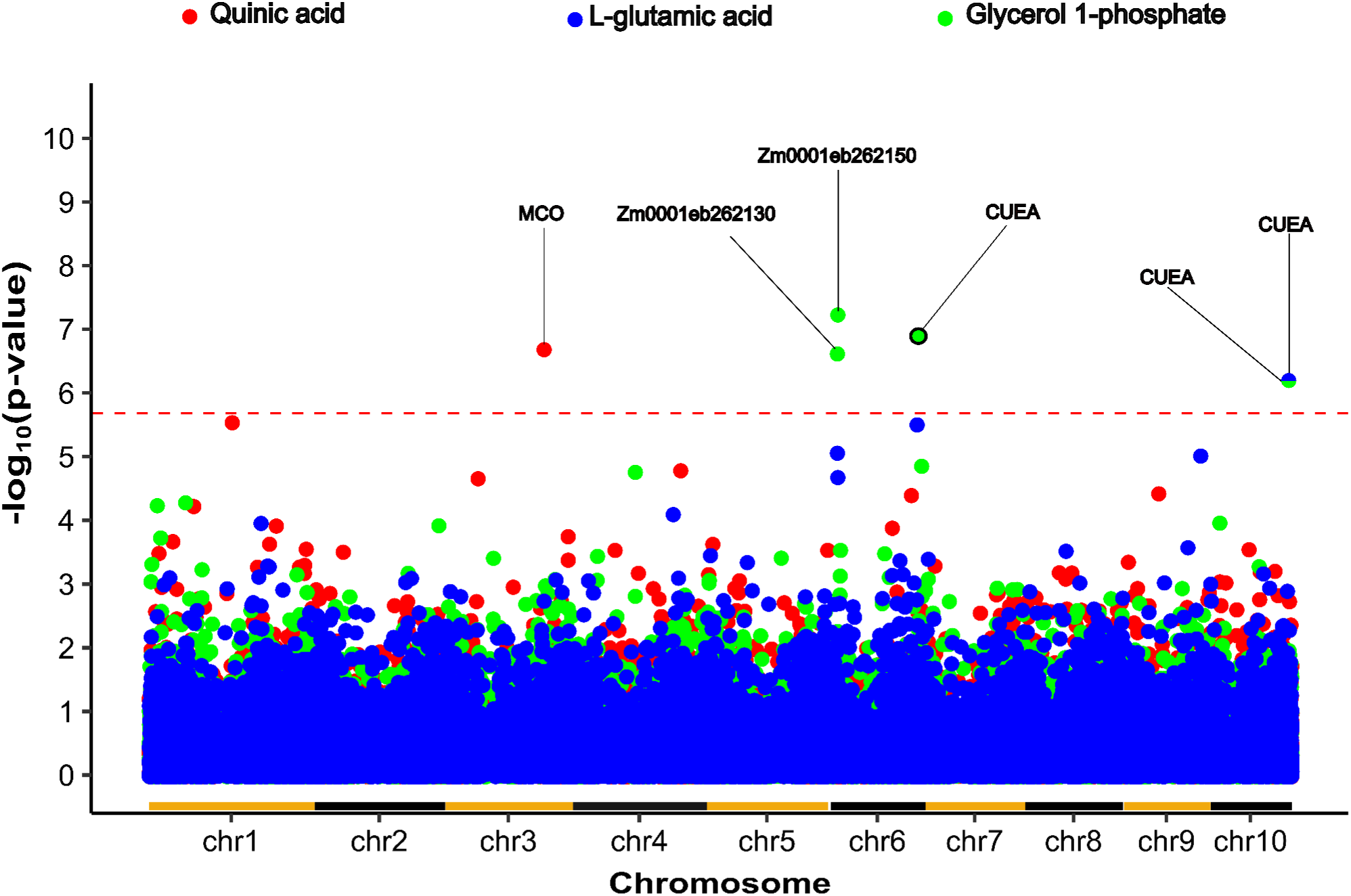
Transcripts associated with variation in metabolite abundance via transcriptome-wide association. Each circle’s position on the x-axis indicates the annotated location of a gene model on the maize genome and its position indicates the statistical significance of the link between variation in the expression of the primary transcript of that gene model and variation in the abundance of a specific metabolite indicated by the color of the dot. Results are shown only for the three metabolites where the significance of at least one transcript exceeded −log_10_(2.03 × 10^−6^), corresponding to the Bonferroni-corrected p-value of 0.05 after correcting for the 24,585 transcripts tested. This threshold p-value is indicated via a horizontal dashed red line. The names of either the proteins encoded by genes above the threshold, or gene model IDs are labeled. Two separate genes, both encoding Cu(2+)-exporting ATPase (Cu(2+)-exporting ATPase) were associated with variation in the abundance glycerol 1-phosphate, one of which was also associated with variation of L-glutamic acid. Alternating color horizontal lines along the x-axis indicate the start and end of each maize chromosome.

We speculated that the limited number of transcripts associated with variation in the abundance of metabolites could reflect nonlinear associations between transcript abundance and metabolite levels. As a complement to conventional TWAS which assumes linear relationships we adopted an explainable AI/random forest-based method^19^, combined with a permutation-based estimate of expected background associations (Figure S10) to identify those transcripts with the most predictive power to explain the abundance of the three metabolites for which at least one significant transcript association was identified via TWAS. A total of 26, 29, and 24 transcripts were identified which exceeded (FDR ≤ 0.05) for L-glutamic acid, quinic acid, and glycerol 1-phosphate respectively (Figure S11). None of the transcripts identified via this method overlapped with genes located near trait-associated markers identified via GWAS, however, One of the six genes identified via TWAS, CUEA, was also identified in the random forest-based analysis. In addition, one of the genes associated with variation in the abundance of L-glutamic acid via random forest N-acetyl-gamma-glutamyl-phosphate reductase (argC) is known to play a critical role in the arginine biosynthesis pathway using glutamate as a precursor^20–22^.

In three cases the same region of the maize genome was linked to variation in metabolite abundance and one of a set of 41 non-metabolic traits, including 28 whole plant phenotypes, 10 traits extracted from hyperspectral leaf reflectance, and three traits related to photosynthetic parameters. Out of 223 genetic markers associated with variation in these non-metabolite traits at a threshold of RMIP ≥ 0.1 (Figure S12), three were located within 100 kilobases of genetic markers associated with variation in the abundance of specific metabolites (RMIP ≥ 0.1). The first of these three cases was an interval of less than 32 kilobases on chromosome 6 containing markers significantly associated with both variation in the abundance of L-serine in mature leaves and variation in the percent grain fill of ears at harvest. This interval contained a gene (Zm00001eb277100) encoding an aldehyde dehydrogenase (ALDH) an enzyme involved in the detoxification of aldehydes by catalyzing their conversion to carboxylic acids, which plays a role in various metabolic pathways, as well as the response to oxidative stress (Figure 4). The second case involved an interval of approximately 64 kilobases on chromosome 1, where markers were significantly associated with both the variation in the abundance of mucic acid and the variation in the LV9 – latent variable 9 – hyperspectral reflectance derive trait. This interval contained a gene (Zm00001eb009750) encoding a transcriptional regulatory protein carrying a Myb/SANT-Like DNA-Binding Domain (Figure 4). The third case was an interval of about 93 kilobases on chromosome 1 that contained markers significantly associated with both variation in the abundance of chlorogenic acid and variation in the number of branches per tassel. This interval included a gene (Zm00001eb035350) encoding a cyclin-dependent kinase (CDK), which is involved in cell cycle regulation (Figure 4).

**Figure 4.**
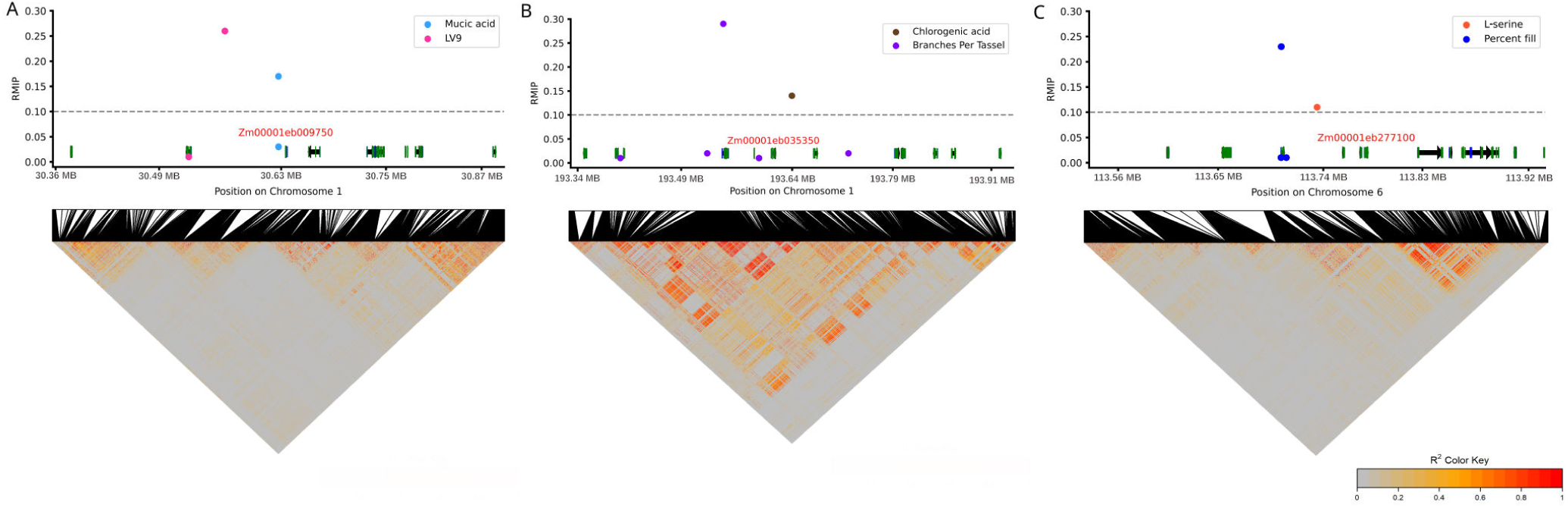
Genomic intervals in maize associated with variation in both metabolic and non-metabolic traits. Genomic intervals that include at least one genetic marker associated with a metabolic trait and at least one genetic marker associated with a nonmetabolic trait (RMIP ≥ 0.1). For each panel, colored points indicate the positions and significance of GWAS hits as shown in Figure 1, black lines indicate the positions of annotated genetic markers within the genomic interval shown, coloration in the triangles at the bottom of the figure indicates the decrease of correlation between different genetic markers in the region, black arrows indicate the positions and strands of annotated genes within the genomic interval shown and green and blue boxes indicate the positions of protein-coding exons and untranslated regions respectively. Candidate genes discussed in the main text or figure legend are labeled in red. **A)** A region (Chromosome 1, 30.35 MB-30.90 MB) containing one marker (chr1:30,626,701) significantly associated with mucic acid abundance and another (chr1:30,562,591) associated with LV9–latent variable 9–a hyperspectral reflectance data derived non-metabolite trait. **B)** A region (Chromosome 1, 193.32 MB-193.95 MB) containing one marker significantly associated with chlorogenic acid abundance and another marker (chr1:193,546,683) associated with the number of branches per tassel. C) A region (Chromosome 6, 113.50 MB-113.94 MB) containing one marker (chr6:113,734,505) significantly associated with L-serine abundance and another marker (chr6:113,703,113) associated with the proportion of the total length of maize ears which develop filled kernels (percent fill).

## Discussion

Understanding the genetic determinants of intraspecies metabolic variation can help to understand causal relationships in plant metabolism and how plant metabolism determines whole plant phenotypes. However, plant metabolism, like transcript abundance, is dynamic and varies throughout the different times and in response to a wide range of environmental signals and perturbations, making it challenging to profile metabolite abundance in comparable conditions from large diversity panels under field-relevant conditions. Here we sought to mitigate the issues of environmental variation and diurnal cycling by employing a set of samples collected in a two-hour period from a large maize diversity panel grown in the field. The patterns of estimated relative abundance for a number of metabolites were reasonably repeatable between independently collected biological samples from the same genotypes (Figure 1A) even before correcting for a number of experimental factors that influenced estimated abundance (Figure 1B). The repeatability was less repeatable than typical whole plant phenotypes scored in the same population (Figure S2; Figure S3; Figure S4).

Some of the lower repeatability of metabolite abundance estimates may be explained by quantification error, patterns of diurnal changes even within the two-hour window of collection, and the high plasticity of plant primary metabolism. However, it should also be kept in mind that typical protocols for scoring many whole plant traits represent averages or aggregate assessments across several (plant height) to dozens (flowering time) of genetically identical plants and these aggregate assessments will tend to reduce residual variance relative to measurements collected from only a single plant. While it remains cost and labor-prohibitive to quantify metabolite abundance in multiple replicated samples from each genotype, as the sampling procedure improves, it may become feasible to collect and pool samples from larger numbers of plants within a single plot, increasing the repeatability of field-measured metabolite abundance.

In any case, genome-wide association studies conducted using dense resequencing-based marker data and the metabolite abundance data generated in this study identified a substantial number of genetic markers that were significantly associated with variation in a number of the metabolites profiled, including 17 marker-trait associations with the highest RMIP scores (all ≥ 0.3). These marker-trait associations were each associated with variation in the abundance of 13 different metabolites (Table 1). In seven cases these markers were located within 50 kilobases of a gene known to play a specific role in plant metabolism, although not necessarily with a role in the expected metabolic pathway. Often windows defined by linkage disequilibrium around individual trait-associated markers included multiple annotated genes including genes of unknown function or possessing only extremely general functional annotations. The ability to link trait-associated genetic markers with a single high-confidence candidate gene remains a major challenge and limitation of genome-wide association studies, even in species such as maize where linkage disequilibrium decays rapidly^23^.

Transcriptome-wide association studies can frequently link specific candidate genes to roles in determining plant phenotypes^16^, even in species with elevated linkage disequilibrium^24^. In our case, we had access to gene expression data generated using paired leaf samples collected from the same plants at the same time as the samples employed for metabolite profiling. In principle, this design should further increase the power for linking transcripts and metabolites as even variation in transcripts induced by non-genetic factors (e.g. diurnal changes, local environmental variation within the field, differences in the leaves selected for sampling by different samples in different plants) could improve the degree of correlation between transcript abundance and downstream metabolic consequences of those same abundance changes. However, despite these advantages, we identified only six significant transcript-metabolite associations via TWAS (Figure 2) and no associations at all for twenty-three of the twenty-six metabolites evaluated. The successful results from GWAS suggest the relatively poor performance of TWAS on the same dataset cannot simply be attributed to the quality/repeatability of the metabolite abundance data. Current best practices for TWAS emphasize focusing on transcripts expressed in more than 50% of the samples examined. This may bias against the discovery of transcripts that exhibit the presence or absence of variation among maize genotypes^25^ and have the potential to play important roles in plant metabolism^26, 27^. In addition, tests for transcript-metabolite associations via transcriptome-wide association studies assume linear relationships between transcript abundance and metabolite abundance. In many causal relationships between transcripts and metabolites, this assumption may not hold true.

We implemented an explainable AI approach based on the random forest algorithm to search for transcripts that exhibit variation in the abundance of specific metabolites in either linear or non-linear fashion^19^. A total of 79 transcripts were linked to the abundance of three metabolites at an FDR threshold of ≤ 0.05, defined based on permutation. Notably, in this analysis, N-acetyl-gamma-glutamyl-phosphate reductase (argC) was linked to variation in the abundance of L-glutamic acid. ArgC catalyzes the reduction of N-acetylglutamate 5-phosphate to N-acetylglutamate 5-semialdehyde in the arginine biosynthesis pathway. This step is critical as it represents a committed stage in the production of arginine. Glutamic acid serves as a precursor in this pathway, and thus its accumulation can be tightly linked to ArgC activity. The proper functioning of ArgC is crucial not only for arginine synthesis but also for overall nitrogen metabolism in plants, impacting growth and stress responses^20–22^.However, the results obtained from this random forest method must be interpreted with some caution. While the method allows the identification of transcripts with nonlinear relationships to metabolites, our current implementation does not rigorously control for the confounding influences of population structure and kinship, which can produce false positives in genome-wide association studies, although the potential for similar effects in transcriptome-wide studies remains less clear.

Of the three methods we employed—GWAS, TWAS, and random forest—GWAS is by far the most generally accepted and widely used. However, we were particularly interested in whether genetic variants that altered plant metabolism would also exhibit impacts on whole plant phenotypes. When a set of 41 non-metabolic traits, including whole plant phenotypes, hyperspectral leaf reflectance, and photosynthetic parameters were analyzed using the same GWAS approach employed for metabolite analysis, three GWAS hits were identified in reasonable proximity to GWAS hits for variation in metabolite abundance. The first case, on chromosome 6, involved a 32-kilobase interval associated with L-serine abundance and percent grain fill, containing a gene encoding aldehyde dehydrogenase. The second case, on chromosome 1, involved a 64-kilobase interval linked to mucic acid abundance and the LV9 trait, containing a gene encoding a Myb/SANT-like DNA-binding domain. The third case, also on chromosome 1, involved a 93-kilobase interval associated with chlorogenic acid abundance and tassel branching, containing a gene encoding cyclin-dependent kinase. The small total number of common genomic intervals identified between metabolite and non-metabolite traits was somewhat unexpected. The use of expanded populations, increased replication, improved protocols for collecting tissue samples, and improved methods for dealing with non-linear interactions may be necessary, either individually or jointly to improve the detection of genetic variants which impact both plant metabolism and non-metabolic traits.

## Materials and Methods

### Field experiments and trait scoring

The maize field experiment from which the plant phenotypes, gene expression data, and metabolite abundance data employed in this study were collected was conducted in the summer of 2020 at the Havelock farm of the University of Nebraska-Lincoln (40.852^◦^N, 96.616^◦^W). The field was laid out in a randomized complete block design on May 6, 2020, consisting of two replications of each genotype. A total of 1680 plots, with each block consisting of 840 plots including 660 entries from the Wisconsin Diversity panel^14^, and the remaining plots consisting of a repeated check line (B97). The layout for each plot consisted of two rows, each 7.5 (about 2.3 meters) feet long, with rows spaced 30 (roughly 0.76 meters) inches apart. Plants within the rows were placed 4.5 (approximately 11.5 centimeters) inches apart from each other, and the plots were separated by 30-inch (around 0.76 meters) alleyways. The experimental design and trait evaluation methodology conducted in the Lincoln, Nebraska field trial has also been previously described^16, 17, 28, 29^.

### Quantification of Metabolite Abundance

On July 8^th^, 2020, when the majority of plots were at the late vegetative or tasseling (VT) stage, duplicate leaf tissue samples were collected from one representative plant per plot in block 1 which consisted of the 840 blocks on the western side of the overall field experiment. Each sample consisted of five leaf disks sampled from the pre-ante-penultimate leaf (the fourth leaf down from the top) of the chosen plant. The leaf tissue was immediately subjected to flash freezing in liquid nitrogen and subsequently stored on dry ice until it could be transferred to a freezer at −80°C. This collection was performed by seven researchers in parallel with all samples collected in two hours of a single day, with collection ending before noon.

One sample per plot was employed for quantification of metabolite abundance. Frozen leaf samples were ground to a fine powder using TissueLyser II (Qiagen). Samples of approximately 25 mg of ground tissue were extracted from each set of ground tissue, precisely weighed, and mixed with 700 *µ*L of methanol and 30 *µ*L of 20 mg/mL ribitol in a 2 ml Eppendorf microfuge by vortexing and stored on ice. Sample tubes were shaken for 15 minutes at 950 rpm on thermomixer at 70°C. Samples were then spun at 17,000 g for 10 minutes and the resulting supernatant was transferred to a new tube and mixed with 325 *µ*L chloroform and 750 *µ*L water by vortexing for 30 seconds. Samples were spun at 1500 g for 15 minutes. Finally, an aliquot of 50 *µ*L from the upper polar phase was transferred into a fresh 2 mL tube and dried with a centrifugal vacuum concentrator. After vacuum drying, each tube was filled with argon gas and tightly closed to prevent the oxidation of metabolites.

Dried metabolite extracts were derivatized by methoxyamination in 20 mg/mL methoxyamine hydrochloride in pyridine for two hours at 37°C. The samples were further trimethylsylilated for 30 minutes at 37°C with 70 *µ*L N-Methyl-N-(trimethylsilyl) trifluoroacetamide (Millipore Sigma). A fatty acid methyl ester mixture was added to the trimethylsylilation solution for retention time calibration. One microliter of each sample was injected into a GC-MS (7200 GC-QTOF system, Agilent) equipped with a HP5msUI (30 m length, 0.25 mm diameter, 0.25 *µ*m thickness) column. GC and MS parameters are exactly as described^30^.

Chromatographic peaks were annotated using MassHunterUnknowns (Agilent) to match retention time and mass spectrum with data in the Fiehn Metabolomics Library (Agilent). Manual curation was used to subset peaks to those which could be confidently identified across all samples in all runs, resulting in a final set of 26 metabolites matched to peaks: aspartic acid, *β* - alanine, caffeic acid, chlorogenic acid, citric acid, glucose, glucose-6-phosphate, fructose, galactonic acid, glyceric acid, glycerol-1-phosphate, glutamic acid, loganin, alanine, threonine, malic acid, mucic acid, myo-inositol, phosphoric acid, quinic acid, raffinose, serine, shikimic acid, sucrose, trans-aconitic acid, and tyrosine. The analysis was conducted over 12 unique batches, with the number of samples per batch varying from 26 to 147, across a total of 27 runs. After subtracting the background noise, the abundance of metabolite was estimated based on the peak height of the representative ion for each metabolite, normalized against the internal standard ribitol, and adjusted for the exact fresh weight of the samples used for extraction. These initial estimates of relative metabolite content were log-transformed prior to downstream analysis.

Initial estimates of metabolite abundance were generated for 795 samples, including 88 observations of B97, the repeated check, and duplicate biological samples collected from different plants in the same plot for 47 additional genotypes. After applying quality control (QC) procedures, which involved removing samples with incomplete data, outliers, or those with inconsistencies in metabolite measurements, data for at least one sample of 660 unique maize genotypes were retained, including 47 genotypes where metabolite abundance was quantified for two duplicate samples collected from separate plants. Although B97 was used as a repeated check, it was not explicitly employed to correct for batch effects. Instead, its inclusion allowed for the assessment of consistency in metabolite measurements across different batches

### Non-metabolite datasets employed in this study

The 28 whole plant phenotypes employed in this study were collected from the same field experiment and the procedure used to measure them, as well as the specific trait values employed are described previously^17^. The ten latent variables employed in this study were also generated from the same field experiment, with hyperspectral leaf reflectance data collected from one leaf per plot using a spectroradiometer^31^ and the resulting data summarized into ten variables through the use of an autoencoder^18^. Fv_P/Fm_P (maximum efficiency of PSII in the light), relative chlorophyll, and leaf temperature were measured in the same field experiment^32^ using MultiSpeq v2 instruments^33^. Gene expression was quantified via RNA-seq from the second set of leaf tissue samples collected in parallel with those employed for metabolite quantification^16^.

### Quantitative Genetic Analyses

Best Linear Unbiased Estimates (BLUES) for each metabolite were estimated using a mixed linear model, generated using the lme4 package^34^ implemented in R v4.2.1^35^ with the equation:

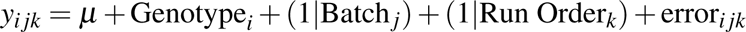

where *y_i jk_* is the mean value for the metabolite of interest in the *i*^th^ genotype, run in the *j*^th^ batch and *k*^th^ run order during the GC-MS pipeline. The variance explained by each factor included in the model was extracted. Repeatability for the estimated abundance of each metabolite was determined using data from 47 genotypes where two independently collected samples were separately processed and quantified. The repeatability was calculated using the following simplified model:

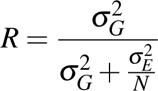

where:

- 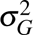 is the genotypic variance, representing the variance explained by genotype.
- 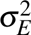 is the residual variance, which includes environmental factors and measurement errors.
- *N* is the number of replicates per genotype (two in this case), adjusting the residual variance accordingly.

Principal component analysis of metabolic abundance data was performed using the `FactoMineR` R package^36^. Prior to conducting GWAS and TWAS, the distribution of BLUEs calculated for each metabolite was manually examined using histograms and scatter plots, and genotypes with extreme values for each metabolites were identified and removed.

### Resampling Model Inclusion Probability Genome-Wide Association

Genome-wide association studies were performed on both metabolite abundance and non-metabolite traits using a set of 660 maize genotypes that had undergone metabolite abundance measurement and passed quality control. A set of 9,794,508 segregating biallelic SNP markers was generated by filtering a larger set of 46 million markers genotyped via resequencing of the Wisconsin Diversity panel^15^ to retain only those with a minor allele frequency greater than 0.05 and heterozygosity less than 0.05 among the 660 maize genotypes included in this study using plink2 (v2.0a1)^37^. The Fixed and Random model Circulating Probability Unification (FarmCPU) algorithm^38^, implemented in the rMVP package^39^, was run 100 times for each phenotype with a different random subset of 10% of phenotypic records masked in each iteration^40^. Four principal components (PCs) calculated from genetic marker data were included in the analysis to control for the confounding effects of population structure. Genetic markers were considered significantly associated with a trait of interest in a given iteration when they exceeded a p-value threshold of 5 × 10^-9^ set using a Bonferroni correction, calculated as 0.05 divided by the total number of SNPs used in the analysis. A given marker-trait association was considered significant if it was identified in at least ten out of the one hundred total GWAS runs conducted, corresponding to a resample model inclusion probability (RMIP) of 0.1.

### Transcriptome-Wide Association Study

TWAS analyses described in this study were conducted using the expression levels measured via RNA-seq^16^ and the compressed mixed linear model^41^ as implemented in the Genomic Association and Prediction Integrated Tool (GAPIT)^42^. For improved comparability, we utilized the same set of 24,585 genes used in the study by Torres et al. (2024)^16^, which were selected based on the following criteria: each gene or transcript that passed the quality filtering process (24,585 gene models with expression ≥ 0.1 TPM in at least 347 of the remaining 693 genotypes) was converted to a range from 0 to 2 using the methodology described by Li et al. (2021)^43^. Briefly, the 5% of samples with the lowest transcripts per million (TPM) values for each gene were scored as 0, the 5% of samples with the highest TPM values for each gene were scored as 2, and the remaining 90% of samples were re-scaled between 0 and 2 using the formula:

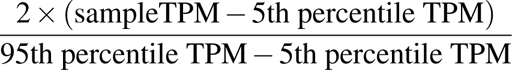

These data were generated using 693 maize genotypes constituting a superset of the 660 maize genotypes employed in this study. The first three principal components of variation calculated by GAPIT from the expression data were included as covariates. Additionally, a kinship matrix was calculated using the VanRaden method^44^, which was used to control for the relatedness among genotypes. A gene was considered significantly associated with the trait of interest when the associated p-value was less than 2.03 × 10^−6^, corresponding to a Bonferroni corrected p-value of 0.05, considering the number of expressed genes employed in the TWAS analysis.

### Random forest-based explainable AI

The random forest algorithm^45^ was employed to predict the genes associated with the metabolites in 660 genotypes using the same gene expression data used for transcriptome-wide association studies. The *randomForest* package in R^46^ was used to build the random forest models with five different tree counts (100, 200, 300, 400, and 500). The *caret* package in R^47^ was then utilized to facilitate 5-fold cross-validation, evaluate model performance using root mean square error (RMSE), and calculate feature importance based on the increase in mean squared error (IncMSE) when a gene was excluded from the model. Twenty control sets were created by shuffling the taxa order while keeping other variables constant. Models were trained, and feature importance scores were calculated using both the original and shuffled datasets. For each metabolite, a threshold corresponding to a false discovery rate of approximately 0.05 was selected based on a comparison of the feature importance scores reported for shuffled datasets.

## Data Availability

The data utilized in this study, including metabolite data for 26 metabolites (Supplementary Table 1), markers associated with 26 metabolite traits and 41 non-metabolite traits identified by GWAS (Supplementary Table 2), and genes associated with three specific metabolites identified by Random Forest (Supplementary Table 3), is publicly available at https://doi.org/10.6084/m9.figshare.26543479. The phenotypic data (Whole plant phenotypes) used in this study was obtained from Mural et al. (2022)^17^ specifically from Supplementary Tables S2 and S3.The hyperspectral used in this study were obtained from Tross et al. (2024)^18^, available at https://doi.org/10.6084/m9.figshare.24808491.v4.Photosynthetic trait data were described and are available from^32^. Estimates of gene expression used in this study are available as part of the supplementary data from Torres et al. (2024)^16^, see associated Figshare repository:https://doi.org/10.6084/m9.figshare.24470758.v1. The genetic marker data employed in this study was generated by filtering the file “WiDiv.vcf.gz” described in^15^ and available from Dryad https://doi.org/10.5061/dryad.bnzs7h4f1 using the criteria described in the methods section.

## Acknowledgements

This project was supported by the Department of Energy under Award Nos. DE-AR0001367 and DE-SC0023138, the Foundation for Food and Agriculture Research under Grant No. 602757, and the National Science Foundation under Grant No. IOS-2332611, and US Department of Agriculture - National Institute of Food and Agriculture under the AI Institute: for Resilient Agriculture, Award No. 2021-67021-35329.

## Author contributions statement

James C. Schnable, Toshihiro Obata, and Ravi V. Mural conceived the experiments, and Connor Pederson and Jonathan Turkus conducted experiments and generated data. Ramesh Kanna Mathivanan, J.Vladimir Torres-Rodriguez, and Nikee Shreshtha analyzed the data and interpreted the results. Ramesh Kanna Mathivanan generated the figures. Ramesh Kanna Mathivanan drafted the manuscript with input from J.Vladimir Torres-Rodriguez, Toshihiro Obata, and James C. Schnable. All authors contributed to the editing of the final manuscript.

## Additional information

James C. Schnable has equity interests in Data2Bio, LLC; Dryland Genetics LLC; and EnGeniousAg LLC. The authors declare no other competing interests.

---

**Figure S1.**
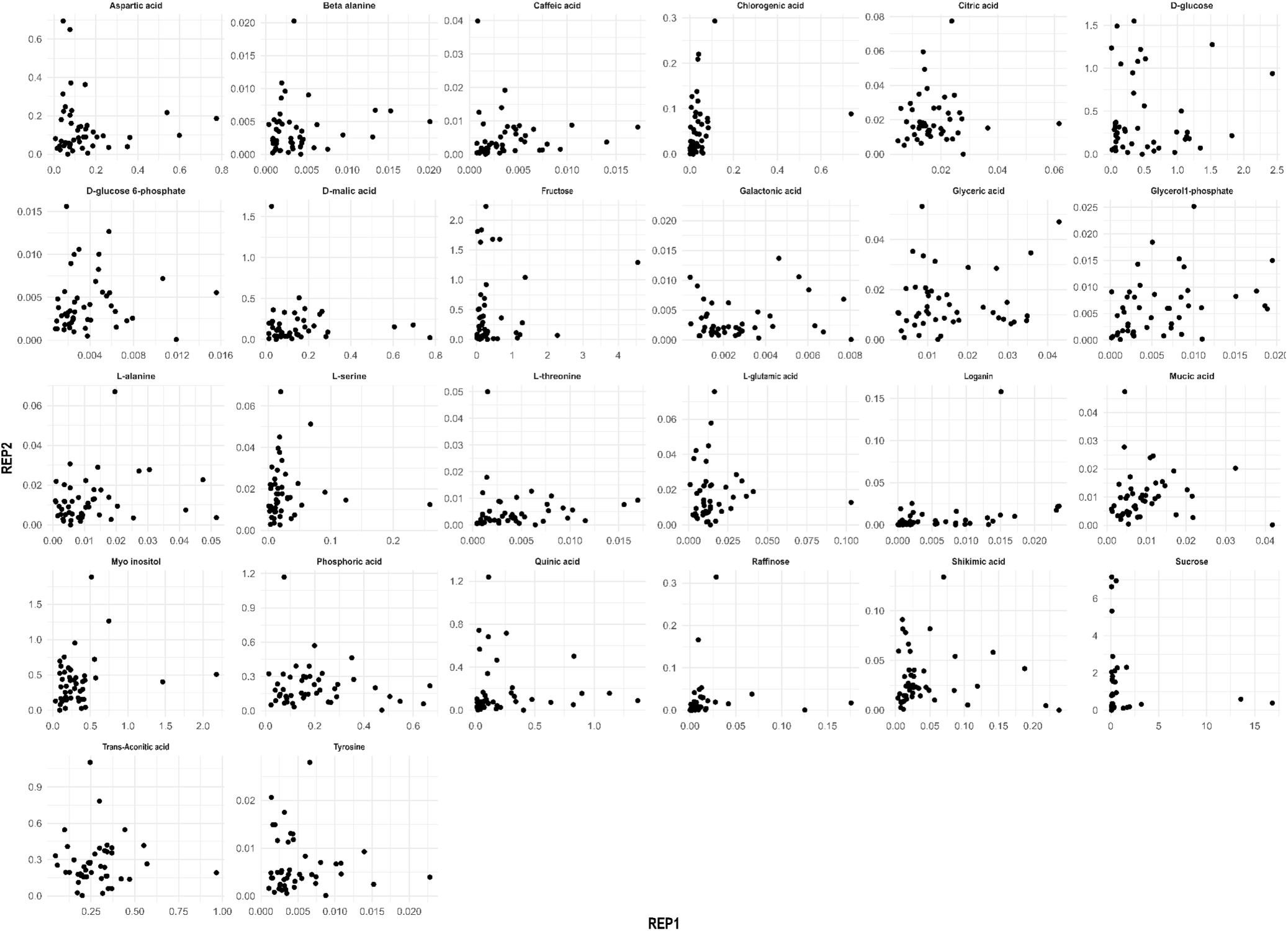
Correlation between 26 metabolite abundance in two replicates of 47 maize genotypes grown in the same field 26 metabolites quantified in a maize diversity panel. REP1 and REP2 correspond to two replications. Metabolites were quantified in two replicates of 47 genotypes.

**Figure S2.**
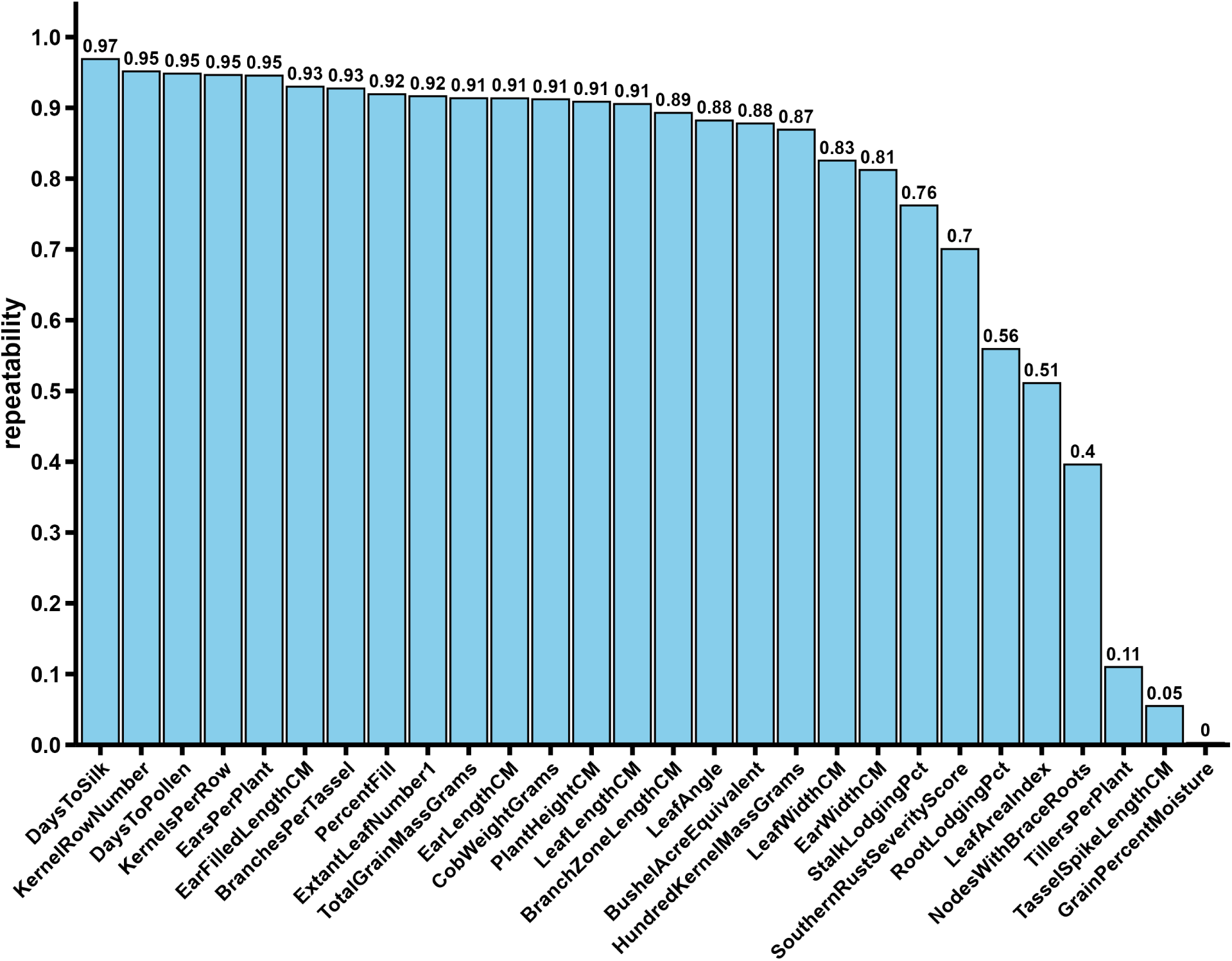
Estimated repeatability of 28 whole plant phenotypes used in this study. Repeatability is defined as the proportion of total variance in metabolite abundance which can be explained by genotype in a dataset of 47 maize genotypes sampled and analyzed twice independently from different plants in the same field.

**Figure S3.**
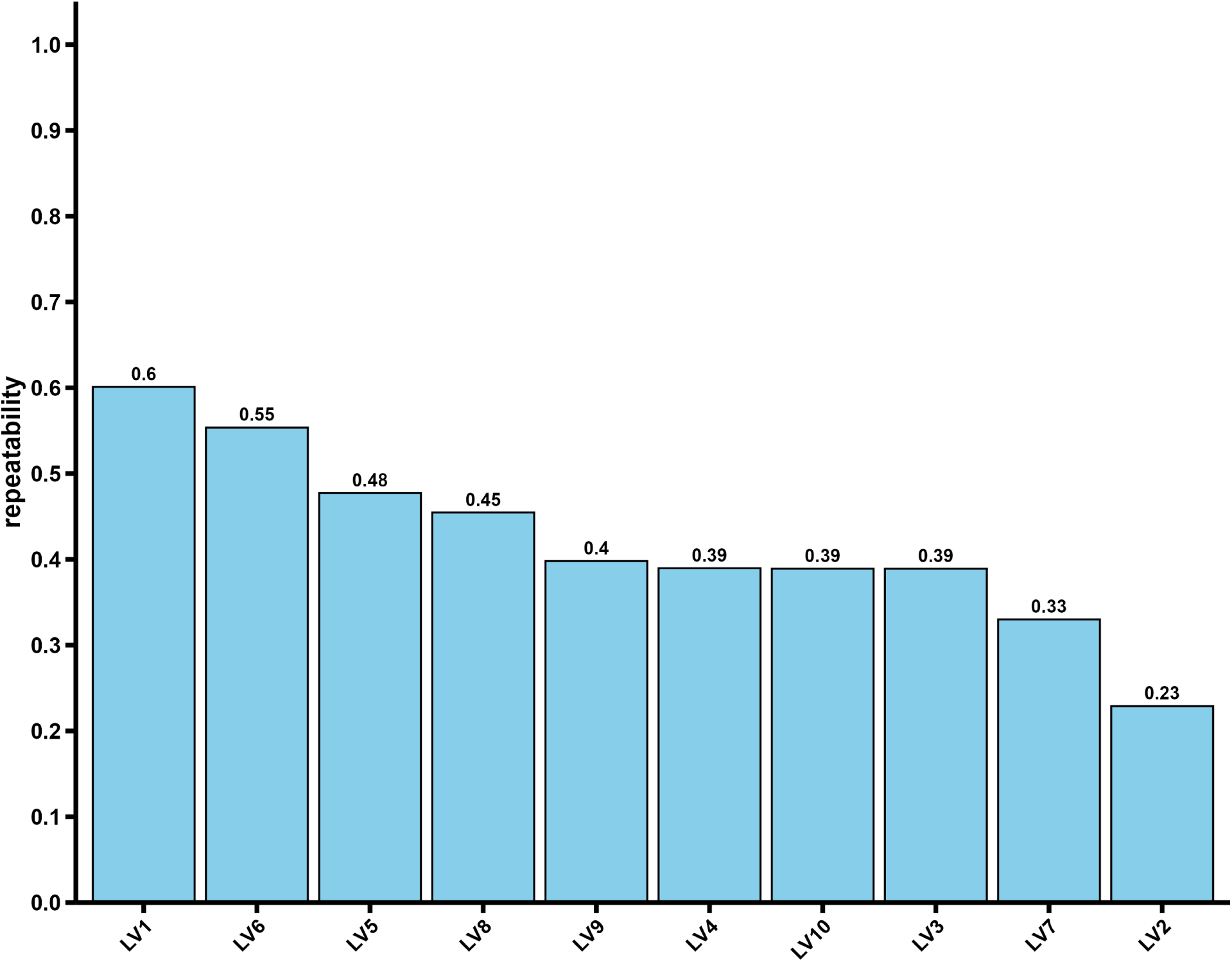
Estimated repeatability of ten hyperspectral leaf reflectance derived latent variables used in this study. Repeatability is defined as the proportion of total variance in metabolite abundance which can be explained by genotype in a dataset of 47 maize genotypes sampled and analyzed twice independently from different plants in the same field.

**Figure S4.**
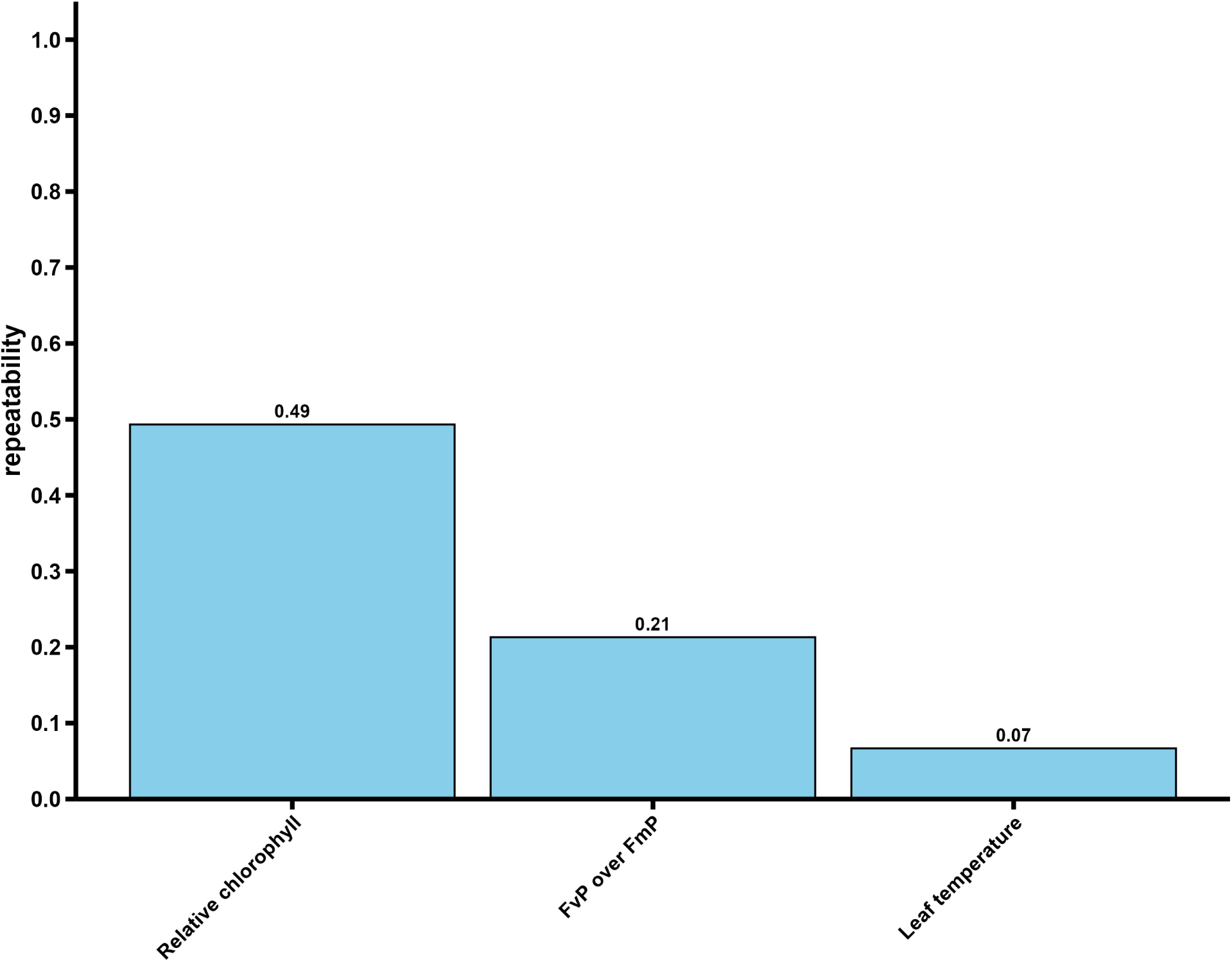
Estimated repeatability of three photosynthesis-related traits used in this study. Repeatability is defined as the proportion of total variance in metabolite abundance which can be explained by genotype in a dataset of 47 maize genotypes sampled and analyzed twice independently from different plants in the same field.

**Figure S5.**
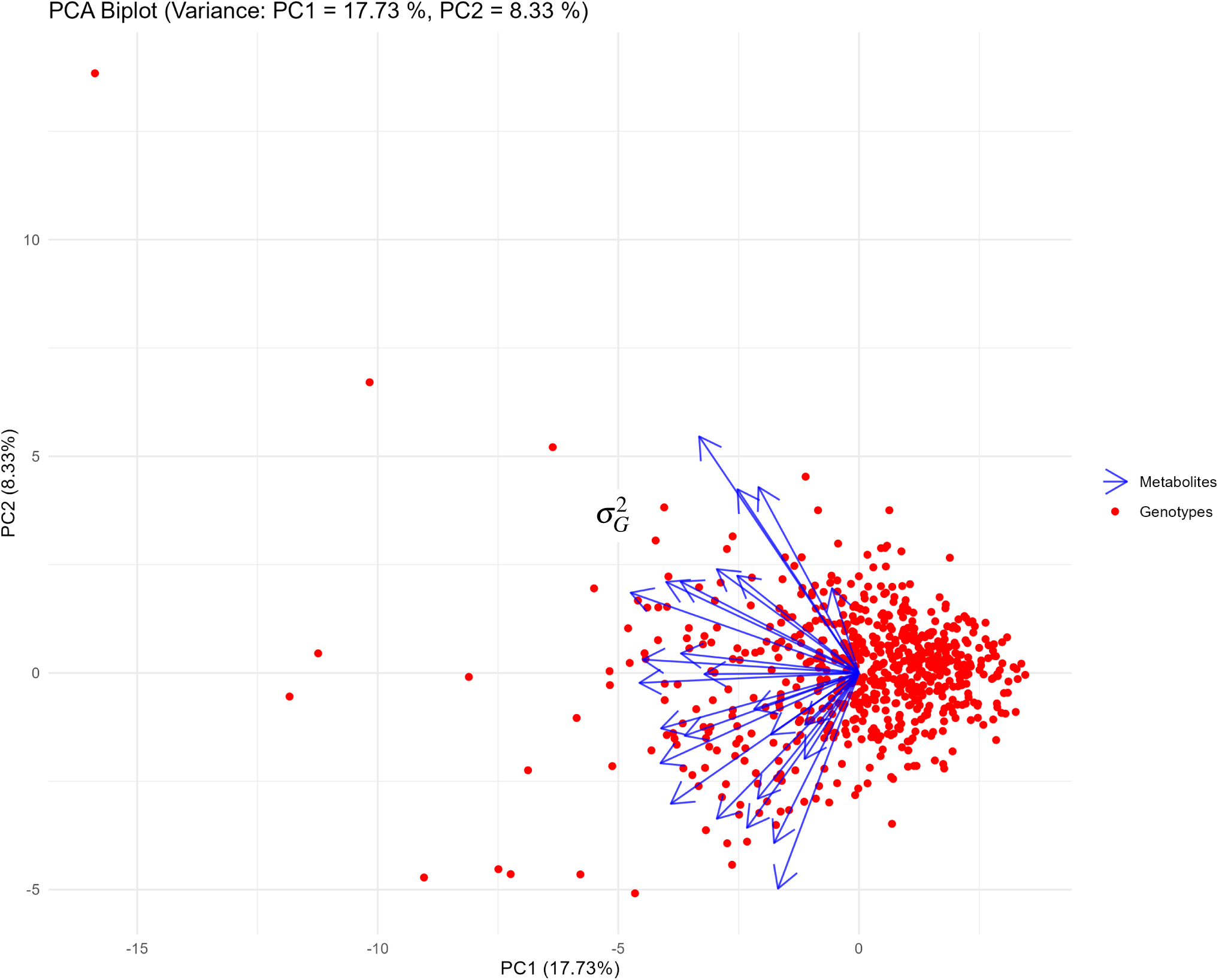
Distribution of scores for the first two principal components of variation in metabolite abundance of a maize diversity panel (n = 795 samples) These blue arrows show the loadings of each metabolite, illustrating their influence on the principal components.

**Figure S6.**
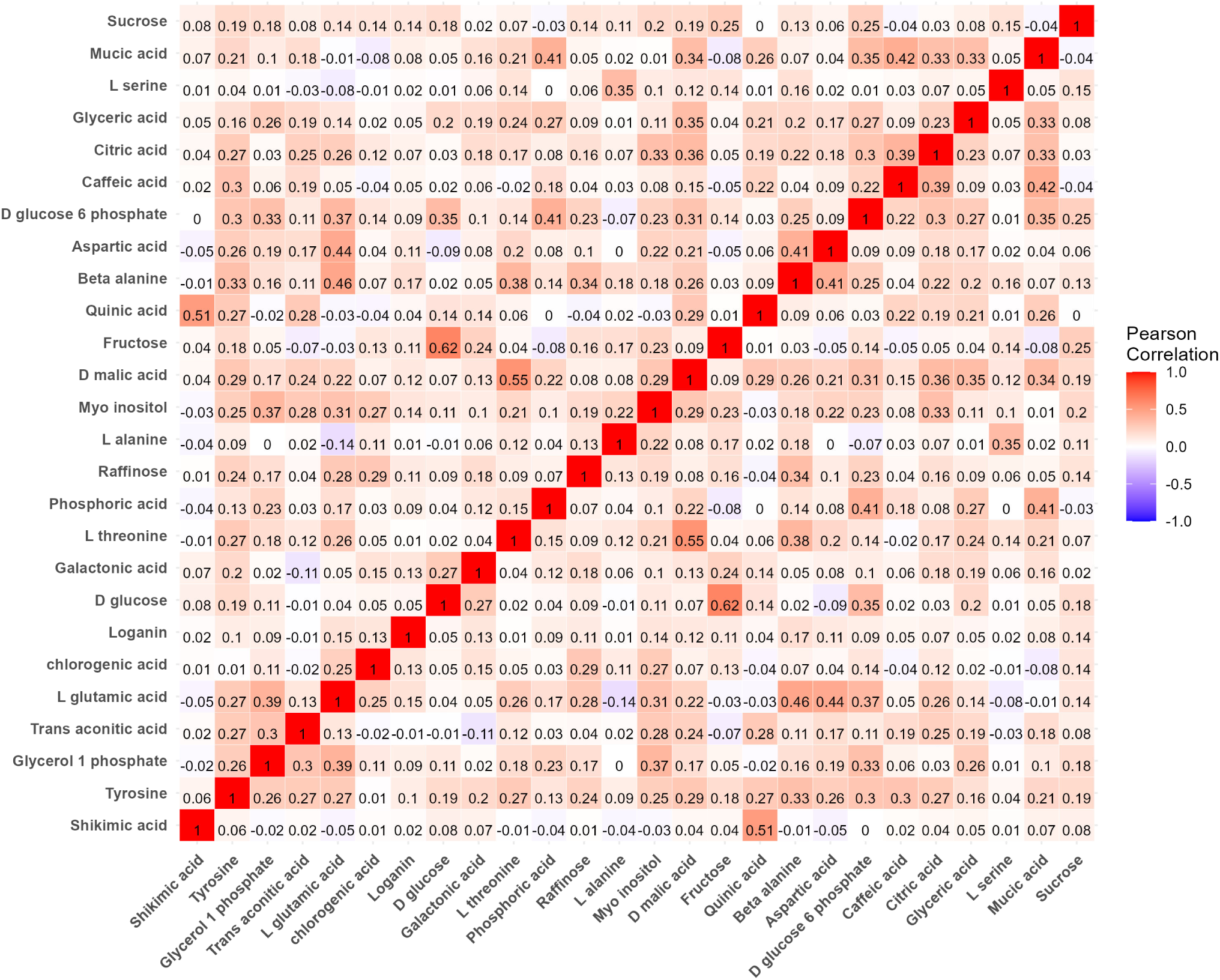
Correlation between the variation of 26 metabolite abundance quantified in a maize diversity panel (n = 795 samples) Dark red squares indicate a strong positive correlation, while dark blue squares indicate a strong negative correlation and lighter colors suggest weaker correlations.

**Figure S7.**
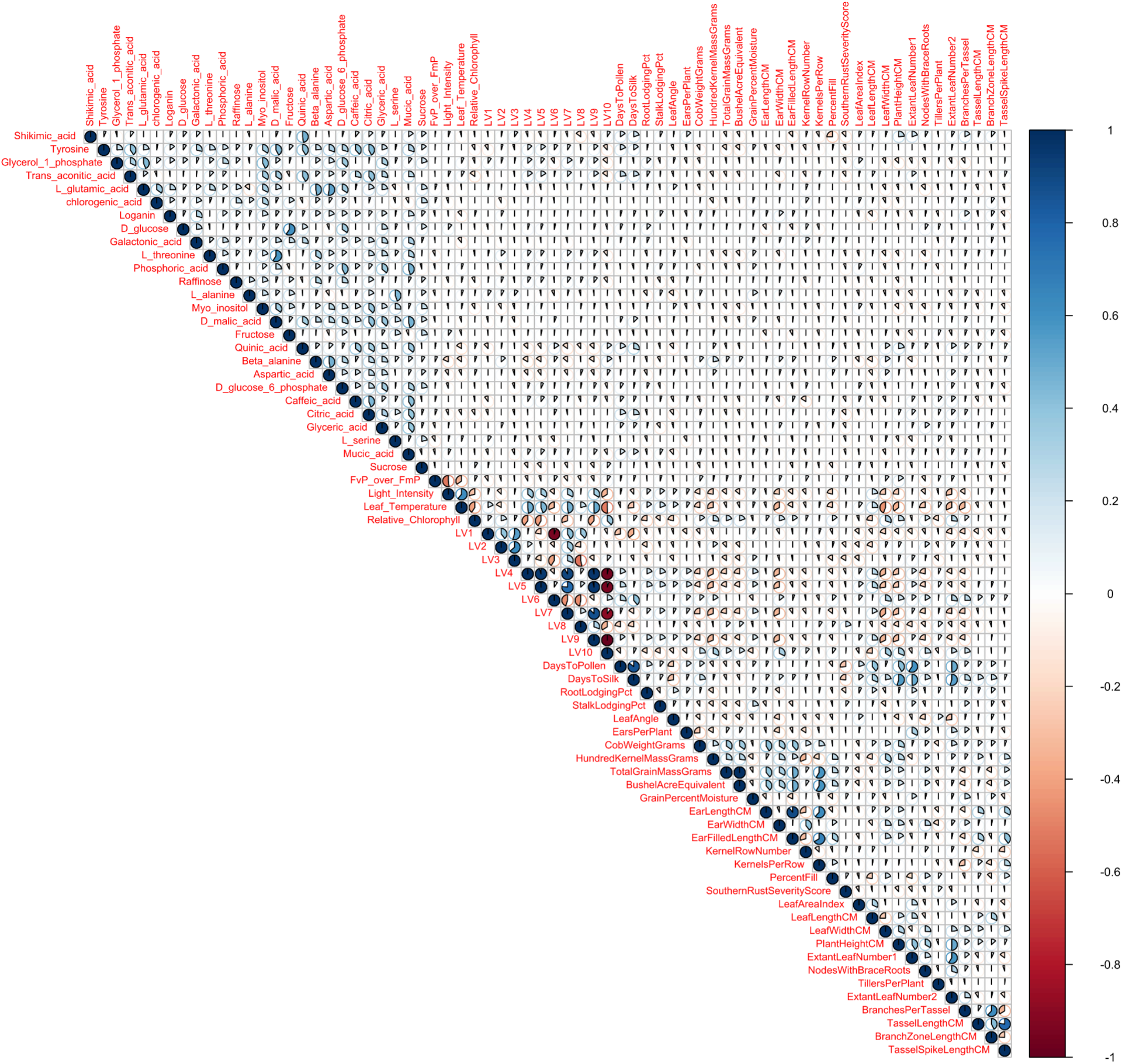
Correlation between the variation of 26 metabolite abundance and 41 non-metabolite traits quantified in a maize diversity panel (n = 795 samples) Dark red squares indicate a strong positive correlation, while dark blue squares indicate a strong negative correlation and lighter colors suggest weaker correlations.

**Figure S8.**
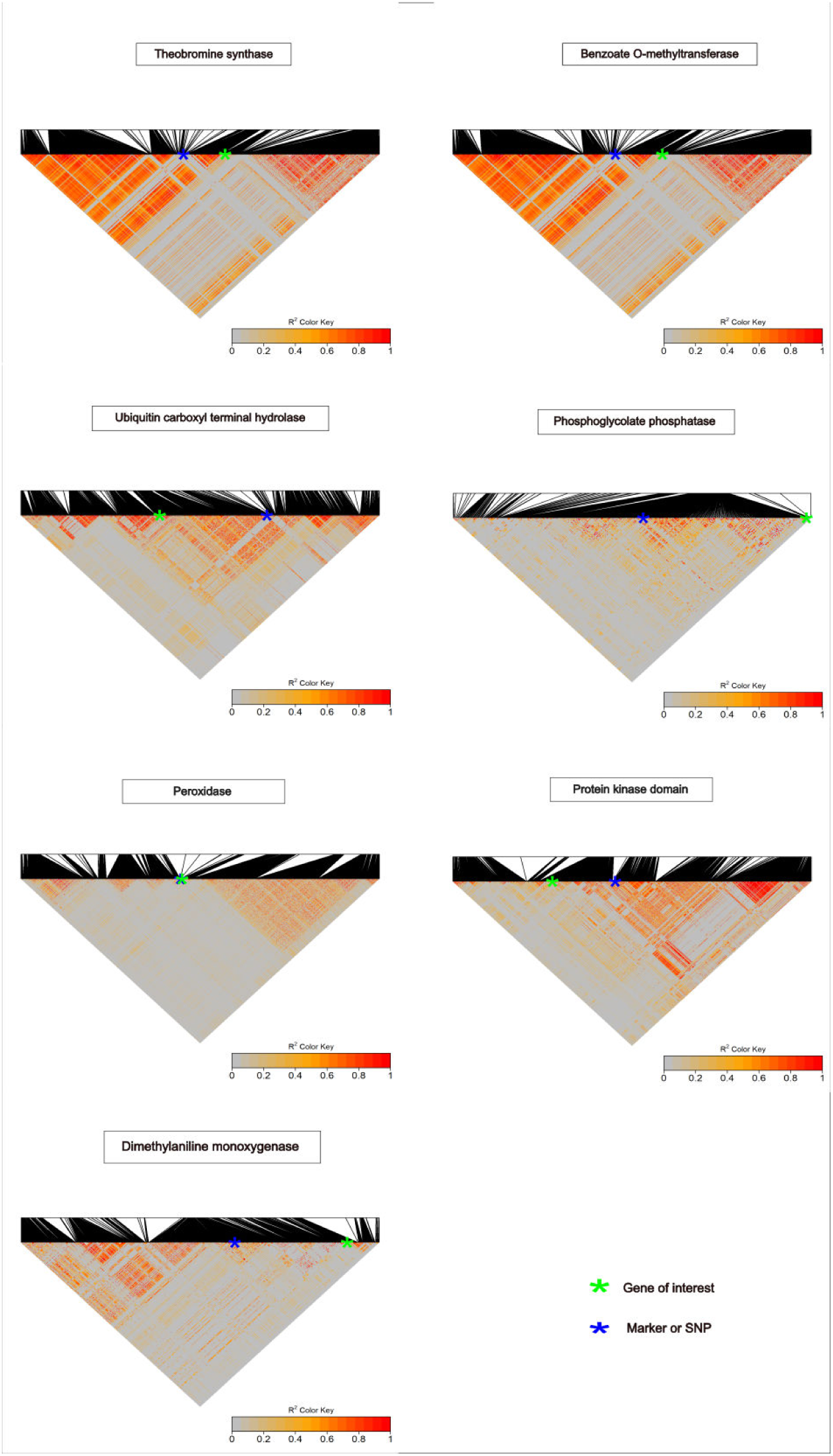
Linkage Disequilibrium (LD) heatmap for seven significant trait-associated SNPs highlighted with closest gene models in Figure 2. The green cross marks the genomic position of the candidate gene model and the blue cross marks the genomic position of trait-associated SNP shown in Figure 2.

**Figure S9.**
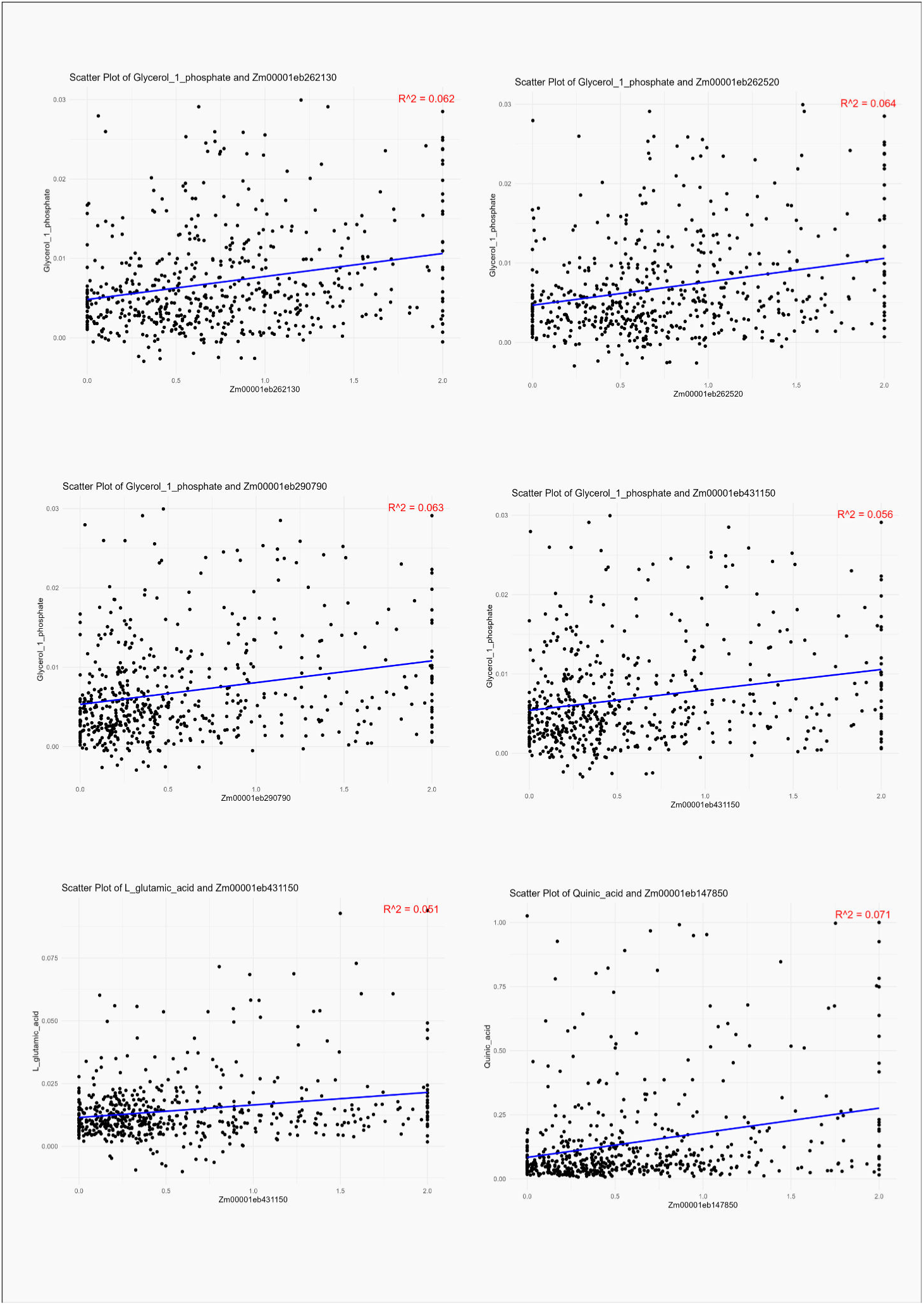
Correlation between three metabolite abundance and gene expression for significant genes identified via transcriptome-wide association studies in Figure 3.

**Figure S10.**
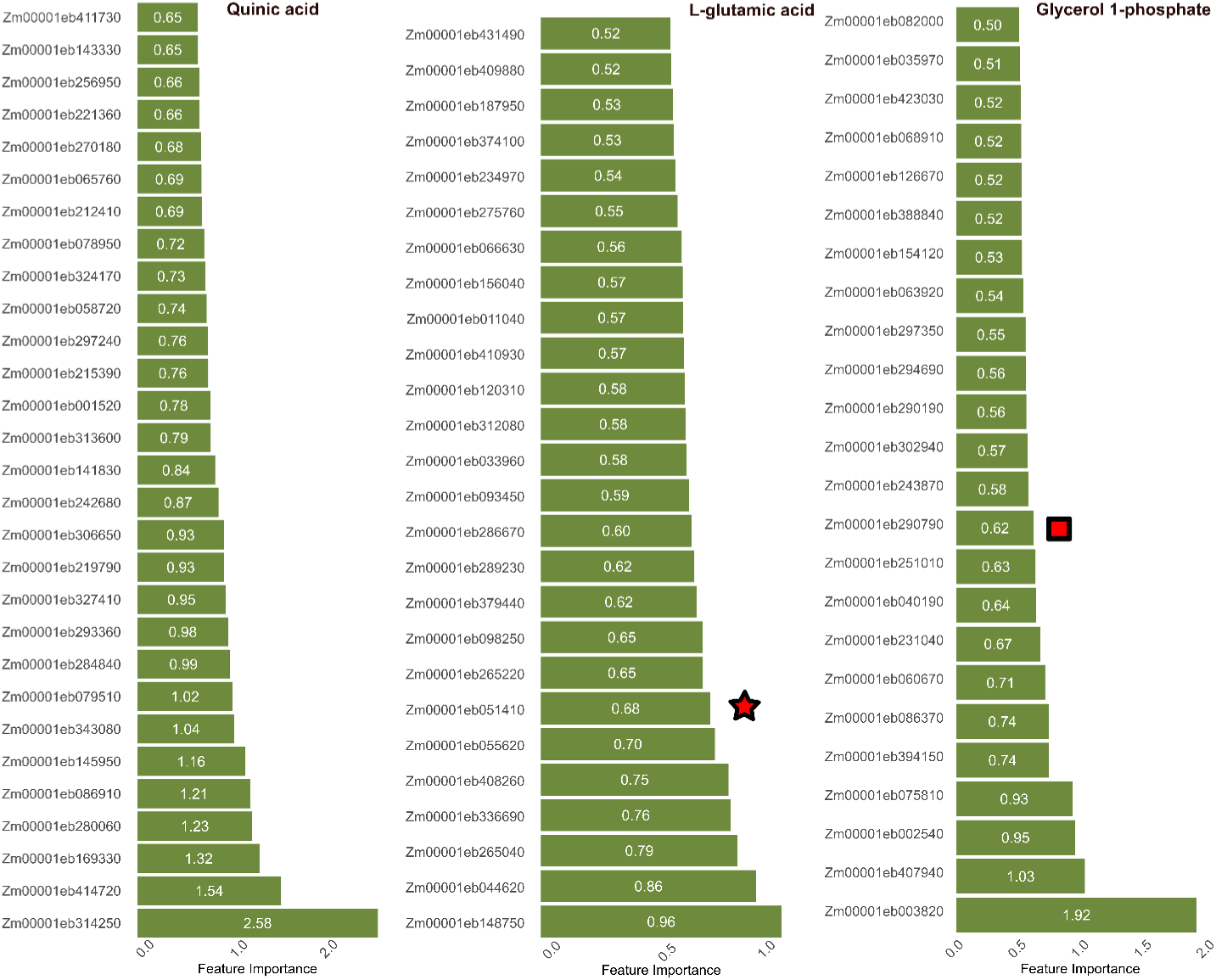
Genes identified with higher feature importance in the random forest (RF) regressor model to predict the abundance of three metabolites in a maize diversity panel. The RF model was trained to predict the abundance of three metabolites out of 26 metabolites in the study based on the significant gene expression-trait associations identified in Figure 3. The number of gene models was selected with higher feature importance than a threshold that correspond to a false discovery rate of approximately 0.05 based on a comparison of the features important scores reported for shuffled datasets for each metabolite. The numbers inside the displayed bar charts represent the feature importance assigned to the gene model for the prediction of corresponding metabolite abundance. The star and square symbol beside gene models indicate that the gene model was found to be significantly associated with the metabolite abundance variation in TWAS analysis.

**Figure S11.**
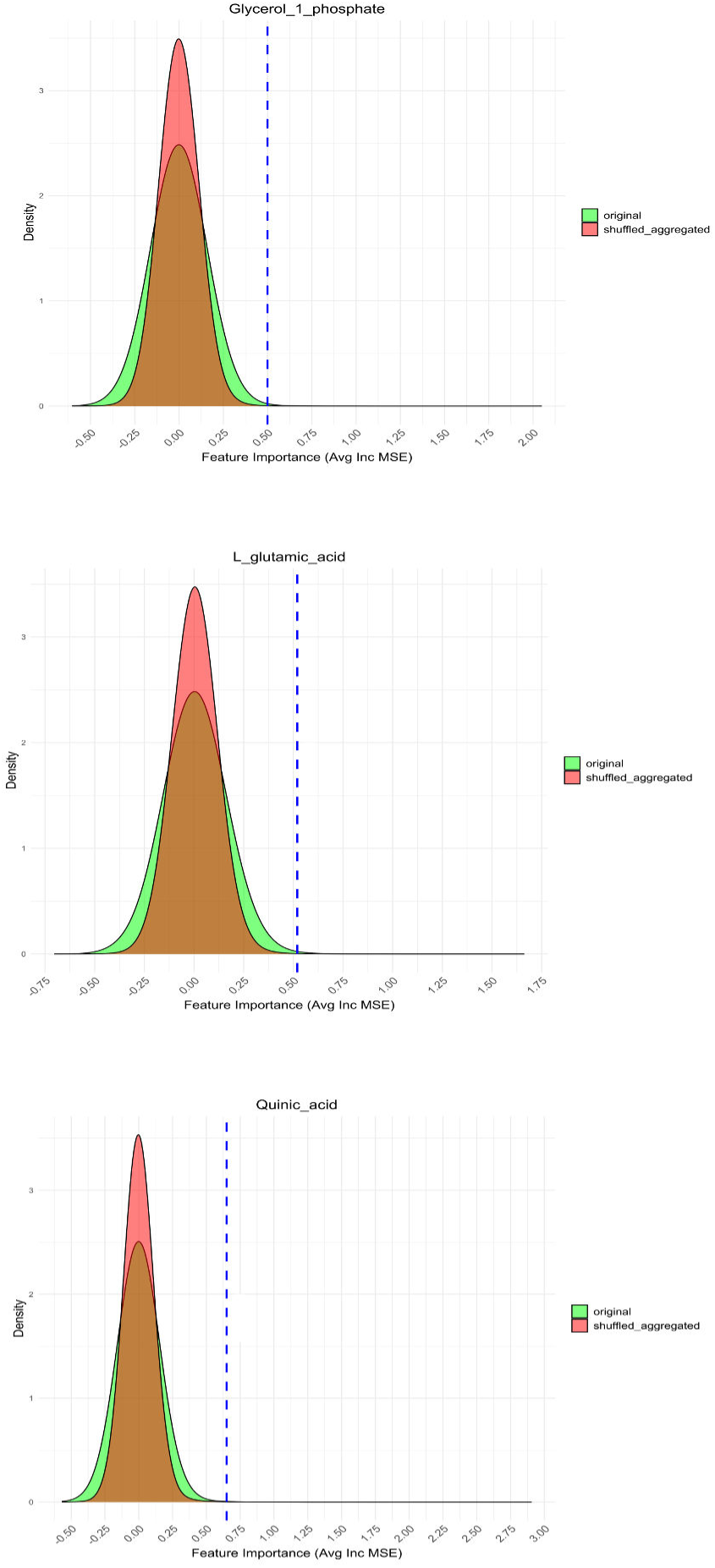
Feature importance scores from original and shuffled data from random forest regressor model using gene expression to predict three metabolites abundance in a maize diversity panel. The blue dashed vertical lines indicate the established threshold for significant feature importance. Genes with feature importance higher than the blue threshold in the original data are considered biologically significant to be associated with metabolite abundance.

**Figure S12.**
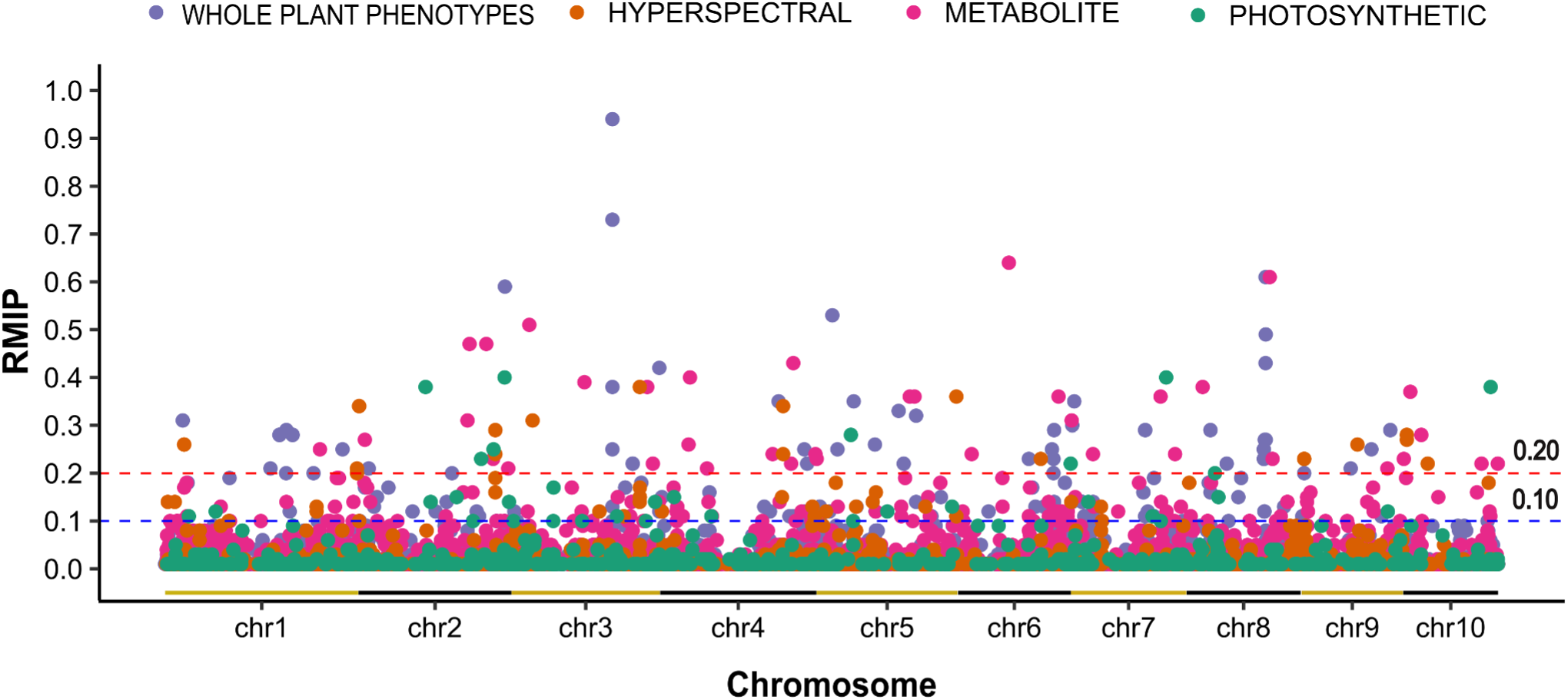
Genetic markers associated with 26 metabolites abundances, 28 whole plant phenotypes, 10 latent variables, and 3 photosynthesis variations via resampling model inclusion probability genome-wide association. Each circle’s position in the x-axis indicates the position of a given genetic marker on the maize genome, and its position on the y-axis indicates the proportion of resampling runs in which the marker was significantly associated with variation in the trait of interest via FarmCPU GWAS. The plot includes two horizontal dashed lines marking RMIP significance thresholds: the upper red dashed line at 0.20 (indicating SNPs significant in at least 20 out of 100 FarmCPU GWAS) and the lower blue dashed line at 0.10 (indicating SNPs significant in at least 10 out of 100 FarmCPU GWAS). Alternating color horizontal lines along the x-axis indicate the start and end of each maize chromosome.

